# *GenomicDataCommons:* a Bioconductor Interface to the NCI Genomic Data Commons

**DOI:** 10.1101/117200

**Authors:** Martin T. Morgan, Sean R. Davis

## Abstract

The National Cancer Institute (NCI) Genomic Data Commons (Grossman et al. 2016, https://gdc.cancer.gov/) provides the cancer research community with an open and unified repository for sharing and accessing data across numerous cancer studies and projects via a high-performance data transfer and query infrastructure. The *Bioconductor* project (Huber et al. 2015) is an open source and open development software project built on the *R* statistical programming environment (R Core Team 2016). A major goal of the *Bioconductor* project is to facilitate the use, analysis, and comprehension of genomic data. The *GenomicDataCommons Bioconductor* package provides basic infrastructure for querying, accessing, and mining genomic datasets available from the GDC. We expect that *Bioconductor* developer and bioinformatics community will build on the *GenomicDataCommons* package to add higher-level functionality and expose cancer genomics data to many state-of-the-art bioinformatics methods available in *Bioconductor*.

**Availability:** https://bioconductor.org/packages/GenomicDataCommons & https://github.com/seandavi/GenomicDataCommons.

## 1 Introduction

Basic and translational cancer research projects – from large, multicenter consortia to individual labs – now produce vast quantities of genomic data. Such datasets are ideally accompanied by extensive phenotypic and clinical information and valuable experimental metadata. To address the needs of the cancer community to access, query, and utilize these cancer-related data resources, the NCI has developed the Genomic Data Commons (GDC, Grossman et al. 2016). The GDC is similar to other large-scale genomics data repositories, while in addtion providing data that have been ‘harmonized’ using stardard, publicly-available pipelines^1^. An additional and unusual feature of the GDC is that users can deposit and then apply standard GDC pipelines to their own genomic data.

The *Bioconductor* project is an open source and open development software project for the analysis and comprehension of genomic and molecular biology data (Huber et al. 2015). Based in the statistical programming language, *R* (R Core Team 2016), the project comprises 1296 interoperable software packages contributed by a diverse community of scientists. Several *Bioconductor* provide access to cancer genomics data (e.g., *RTCGA* (Kosinski and Biecek 2016), *TCGABiolinks* (Colaprico et al. 2016), *RTCGAToolbox* (Samur 2014)) but they are focused, primarily, on TCGA data and, in some cases, rely on legacy data sharing platforms. The *GenomicDataCommons* software package provides infrastructure for finding, accessing, and downloading cancer genomics data from the NCI GDC. Our expectation is that the *GenomicDataCommons* package will facilitate use of GDC data by the larger *Bioconductor* community and increase the effiency of drawing biologically important conclusions from published cancer genomic data.

## 2 Design and Usage

The goal of the *GenomicDataCommons Bioconductor* package is to expose the GDC RESTful application programming interface (API^2^) to *Bioconductor* users via an *R* client interface. The *GenomicDataCommons* package supports both the metadata query and data download capabilities of the GDC API. The *GenomicDataCommons* package is implemented as a set of S3 classes and methods with accompanying utility functions.

### 2.1 Usage

Finding data in the GDC starts with constructing a query of available metadata. We model our *R* interface for querying GDC metadata on a common approach, using pipes to connect functional methods. This is similar to the approach used by popular packages such as *dplyr* (Wickham and Francois 2016). Querying the GDC API to get the unique file ids of all HTSeq-quantified gene expression data from the TCGA-GBM project forms an illustrative example:

~~~
library(GenomicDataCommons)
file_records = files() %>%
     filter(^˜^ cases.project.project_id == 'TCGA-GBM' &
     data_type == 'Gene Expression Quantification' &
     analysis.workflow_type == 'HTSeq - Counts'
  ) %>%
     select(default_fields('files')) %>%
     response_all()
~~~

The `files()` call creates a `GDCquery`. The `filter()` verb allows *R*-like syntax to generate filter criteria (that are then translated to an *R* list); note that field names like “data_type” can be used as *R* names using ‘non-standard’ evaluation. The `select()` verb then sets the return fields, which, in this case are limited to the GDC-specified `default_fields`. Up to this point, no data have been retrieved from the GDC. The `response_all()` call performs the actual metadata retrieval, returning a `GDCResponse` object.

The GDC API also supports simple aggregation based on metadata fields, such as the number of cases summarized by GDC project.

~~~
cases_by_project = cases() %>%
      facet('project.project_id') %>%
      aggregations()
head(cases_by_project)
~~~

### 2.2 Downloading data

Downloading single files or multiple small files is supported directly from *R*. For larger sets of files or single large files, we use the GDC data transfer tool and a ‘manifest’ file. As an example, to create a manifest from the file records collected above, we use the `manifest()` verb.

~~~
manifest_df = file_records %>% manifest()
~~~

The `transfer()` function will use the GDC transfer client (a separate command-line tool) to perform robust, performant download of the data.

### 2.3 BAM file slicing

The BAM file is the current standard for storing reads aligned against a reference. Many use cases for these alignments require only a particular genomic region rather than all alignments. BAM ‘slicing’ is implemented as a basic wrapper upon which developers can build functionality.

~~~
# *get 10 RNA-seq BAM files*
file_ids = files() %>%
    filter(
        ^∼^ data_format=='BAM' & experimental_strategy=='RNA-Seq'
    ) %>% response() %>% ids()
~~~

The file ids, all associated with aligned BAM files, can be used for region-based slicing.

~~~
# *An access token from the GDC is needed here*
# *for the the controlled-access BAM files*.
bamfile = slicing(
      file_ids[1], regions = 'chr17:74000000-76000000', token=gdc_token()
)
~~~

The resulting BAM file, with the file name `bamfile`, can be accessed using any of the standard *Bioconductor* tools for working with aligned reads in BAM format (see, e.g., *GenomicAlignments*).

## 3 Conclusions

The GDC is the latest iteration of a genomic data repository from the NCI. The GDC offers excellent web-based query interfaces and a growing number of web-based tools for interacting with data. The *GenomicDataCommons* package provides bioinformatics and genomic data scientists with programmatic tools that enable statistical analysis, version control, and principles of reproducible computational research (Sandve et al. 2013). The availability of *GenomicDataCommons* in *Bioconductor* proides a large, innovative community of researchers and software developers, many of whom focus on cancer research, ready access to the rich data resources of the GDC. The reproducibility, interoperability, usability, sustainability, and continuous integration that are intrinsic to *Bioconductor*, combined with a plethora of existing tools, have produced a multiplier effect for software contributed to the project, making it an excellent environment for cancer genomic data science. The *GenomicDataCommons* package forms the foundation to build and rapidly iterate on innovative tools for the analysis and comprehension of cancer genomics datasets available at the NCI GDC.

## Footnotes

1 See https://docs.gdc.cancer.gov/Data/Introduction/ for details.

2 https://docs.gdc.cancer.gov/API/Users_Guide/Getting_Started/

